# Temporal dynamics and representational consequences of the control of processing conflict between visual working memory and visual perception

**DOI:** 10.1101/2023.12.07.570647

**Authors:** Chunyue Teng, Jacqueline M. Fulvio, Mattia Pietrelli, Jiefeng Jiang, Bradley R. Postle

**Author notes:** Correspondence should be addressed to: Chunyue Teng, Department of Neuroscience Lawrence University Appleton, WI 54911, USA.

## Abstract

Visual working memory (WM) extensively interacts with visual perception. When information between the two processes is in conflict, cognitive control can be recruited to effectively mitigate the resultant interference. The current study investigated the neural bases of the control of conflict between visual WM and visual perception. We recorded the electroencephalogram (EEG) from 25 human subjects (13 male) performing a dual task combining visual WM and tilt discrimination, the latter occurring during the WM delay. The congruity in orientation between the memorandum and the discriminandum was manipulated. Behavioral data were fitted to a reinforcement-learning model of cognitive control to derive trial-wise estimates of demand for proactive and reactive control, which were then used for EEG analyses. The level of proactive control was associated with sustained frontal-midline theta activity preceding trial onset, as well as with the strength of the neural representation of the memorandum. Subsequently, discriminandum onset triggered a control prediction error signal that was reflected in a left frontal positivity. On trials when an incongruent discriminandum was not expected, reactive control that scaled with the prediction error acted to suppress the neural representation of the discriminandum, producing below-baseline decoding of the discriminandum that, in turn, exerted a repulsive serial bias on WM recall on the subsequent trial. These results illustrate the flexible recruitment of two modes of control and how their dynamic interplay acts to mitigate interference between simultaneously processed perceptual and mnemonic representations.

**Significance Statement:** One hallmark of human cognition is the context dependent, flexible control of behavior. Here we studied the “mental juggling” required when, while holding information in mind, we have to respond to something that “pops up in front of us” before returning to the interrupted task. Using parameter estimates from a reinforcement-learning model, we analyzed EEG data to identify neural correlates of two discrete modes of cognitive control that act to minimize interference between perception and WM. Proactive control, indexed by frontal midline theta power, increased prior to trial onset when a high level of conflict was expected. Reactive control acted to suppress the representation of items likely to interfere with performance, a processing step with consequences for the subsequent trial.

## Introduction

Holding task-relevant information in working memory (WM) while simultaneously processing sensory input is essential for successfully carrying out everyday behavior (e.g., Boettcher et al., 2021; Desimone & Duncan, 1995; D’Esposito & Postle, 2015; Baddeley, 1992). This mental juggling act can be challenging when the two types of information are in conflict (e.g., Bae & Luck, 2018; Lorenc et al., 2021; Rademaker et al., 2015; Teng & Kravitz, 2019). Although the interaction between WM and perception may be obligatory (e.g., Soto et al., 2005, 2008; Olivers et al., 2006; Kiyonaga & Egner, 2016; Gayet et al., 2013; Ding et al., 2019; Teng & Kravitz, 2019; Teng & Postle, 2021; Fukuda, et al., 2022; Rademaker et al., 2015), it is possible to mitigate interference between the two via the application of cognitive control (e.g., Teng et al., 2022; Carlisle & Woodman, 2011; Kiyonaga et al., 2012; Kiyonaga & Egner, 2014). Here we seek to elucidate the electroencephalographic (EEG) bases of the control of conflict between WM and visual perception.

Cognitive control has been widely examined in tasks that induce conflict between stimulus-response associations, that require response inhibition, that involve error processing, or that impose a punishment (e.g., Aron, 2007; Botvinick et al., 2001; Egner, 2017). Important for the present study is the dual mechanisms of control framework (Braver, 2012), which posits two distinct modes of cognitive control: proactive and reactive. Proactive control refers to a tonic, sustained state of elevated control (e.g., activation of goal-relevant rules) when conflict is anticipated, whereas reactive control refers to the phasic “after-the-fact” recruitment of control that is recruited in response to an unexpected instance of conflict. These two modes of control can be manipulated independently. In the color-word Stroop task (Stroop, 1935), for example, increasing the proportion of incongruent trials within a block engages proactive control, and consequently reduces the average Stroop interference effect during these blocks (e.g., Bugg, 2012; Braver et al., 2021). Reactive control, in contrast, is evidenced with the manipulation of item-specific proportion congruency: Stroop interference decreases for the frequently incongruent item, suggesting an association between predicted item-level congruity and control demands (e.g., Bugg, 2012; Bugg & Crump, 2012; Gonthier, Braver, & Bugg, 2016).

Neuroimaging studies indicate that proactive control elicits sustained elevated levels of activity in lateral prefrontal cortex (PFC) (e.g., Burgess & Braver, 2010; Braver, 2012), whereas reactive control elicits transient activation of the lateral PFC (e.g., Burgess & Braver, 2010; Feredoes et al., 2006) and other regions including the anterior cingulate cortex (e.g., Brown & Braver, 2005; Chiew & Braver, 2017).

In the present experiment we recorded the electroencephalogram (EEG) while subjects performed a dual-task procedure that combined WM for orientation of a Gabor patch (”memorandum”) with a clockwise-counterclockwise discrimination of a second Gabor patch (“discriminandum”) presented during the WM delay, with the congruity of the two the key experimental manipulation (Figure 1A). Although the WM and discrimination decisions were independent, previous work indicates that they interact: WM recall precision is higher, and visual discrimination faster and more accurate, when memorandum and discriminandum are congruent (Teng et al., 2022). Furthermore, fits of these data to a reinforcement learning-based Flexible Control Model (FCM; Jiang et al., 2014, 2015) have provided evidence for the adaptive adjustment of control: The level of predicted conflict was negatively related to the behavioral cost of memorandum-discriminandum incongruity; and when the expected level of conflict did not match the actual congruity between discriminandum and memorandum, the incongruent discriminandum exerted a repulsive serial bias on WM recall on the subsequent trial (Teng et al., 2022). (Congruent discriminanda exerted the more commonly observed attractive serial bias.) Our goal with the present work was to use fits of the FCM to behavioral data from the WM/discrimination dual task (Teng et al., 2022) to identify and distinguish EEG correlates of proactive and reactive control processes recruited to control WM-perception interactions.

**Figure 1.**
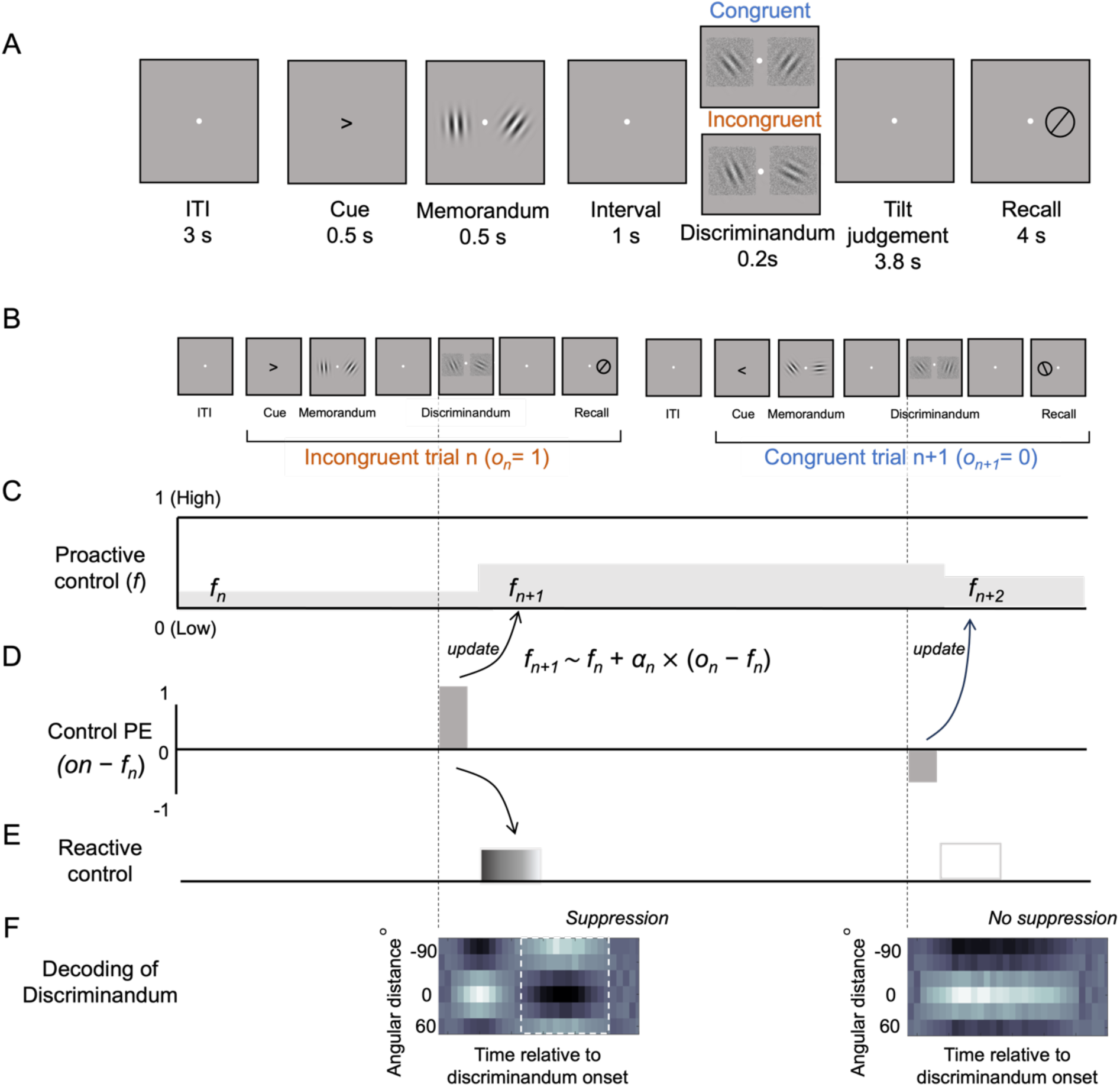
Task structure and the hypothesized interplay between proactive and reactive control. (A) Each trial started with a centrally presented arrow cue that indicated which side of the screen to attend to for both memory and discrimination tasks. Subjects first memorized the orientation of a Gabor patch (”memorandum”), and then during the WM delay, they performed a speeded clockwise/counterclockwise tilt judgment on the noise-covered Gabor patch (“discriminandum”). The orientation of the discriminandum was congruent with that of the memorandum on half of the trials, and incongruent with it on the other half of the trials. (B) Illustration of two consecutive trials of the dual WM+discrimination task, with the orientation of the discriminandum incongruent with the memorandum on the first trial illustrated (“*n”*), and congruent on trial *n+1*. (C): The dynamics of proactive control across the two trials illustrated in (B). In the FCM, proactive control is operationalized by the *predicted conflict* (*f*) parameter, and here it is illustrated with gray shading. For the situation illustrated here, the value of *f* preceding the onset of trial *n* is relatively low, and this persists at this level until it is updated following the onset of that trial’s discriminandum. Because the discriminandum on trial *n* is incongruent with the memorandum, the value of *f* increases, and this results in an increase in the tonic level of proactive control that persists until the onset of the discriminandum on trial *n+1*. (D) The *control PE* on these two trials. On each trial, upon discriminandum onset, a signed *control PE* signals the discrepancy between the level of *f* and the observed congruity (*o_n_ - f_n_*). On trial *n*, incongruity results in a positive *control PE* (because *o_n_* takes a value of 1), whereas on trial *n+1*, congruity results in a negative *control PE* (because *o_n_* takes a value of 0). The *control PE* has two functions. First, it is used to update *f*. Second, it triggers phasic reactive control. (E) The relatively large positive *control PE* on trial *n* triggers a relatively strong recruitment of reactive control, whereas the negative *control PE* on trial *n+1* results in no recruitment reactive control (Note that the effects of the *control PE* are assumed to be task specific: in this task an incongruent discriminandum recruits reactive suppression (so as to minimize interference with recall of the memorandum), but a congruent discriminandum does not require the application of control). (F) Hypothesized decoding results of the discriminandum, plotted as pattern similarity between the test orientation and representational templates from the training set. On trial *n*, reactive control is implemented as suppression of the neural representation of the incongruent discriminandum. On trial *n+1*, in the absence of reactive control, the representation of the discriminandum gradually decays.

## Methods

### Subjects

A total of 25 subjects (12 female and 13 male, mean age = 22.3 ± 4.6) were recruited from the community of the University of Wisconsin–Madison in exchange for monetary compensation. The sample size was determined based on previous studies that investigated similar event-related potential (ERP) signatures of frontal midline theta power in the context of cognitive control (e.g., Cohen, 2016; Messel et al., 2021; Töllner et al., 2017; Whitehead et al., 2019). All participants reported being neurologically healthy, right-handed, and having normal or corrected-to-normal visual acuity. The study was approved by the University of Wisconsin– Madison Health Sciences Institutional Review Board.

### Stimuli and procedure

The stimuli were created and presented using MATLAB (The Mathworks) and the Psychtoolbox extension (Brainard, 1997). They were displayed on a 60 x 34 cm screen with a refresh rate of 60 Hz. Participants viewed the stimuli from a distance of approximately 58 cm in a dimly lit room.

The experiment had a dual-task structure: WM delayed recall (a.k.a. “delayed estimation”) with an independent perceptual discrimination occurring during the WM delay period. The background was gray throughout the experiment. Each trial began with a centrally presented left- or right-pointing arrow (black, 0.8° × 0.8°), its direction indicating the side of screen to attend to for that trial. After 0.5 seconds, two memoranda (Gabor patches: radius = 6°; contrast = 0.8; spatial frequency = 4 cycles/°; phase angle randomized between 0 to 180 degrees; 10° apart) were presented, centered on the horizontal meridian, one on each side of a white central fixation dot, for 0.5 seconds. Subjects were instructed to memorize the orientation of the memorandum that appeared on the cued side of the screen, but to maintain central fixation.

After a delay of 1 second, two discriminanda (Gabor patches, same properties as the memoranda) appeared at the same two locations for 0.2 seconds. Subjects were instructed to judge the tilt of the discriminandum on the cued side relative to vertical, and indicate their response by pressing, with fingers of the left hand, the ’E’ key for a clockwise tilt and the ’F’ key for a counterclockwise tilt. 2.8 seconds after the offset of the discriminanda, a recall dial, displaying a randomly determined orientation, appeared at the location of the cued memorandum, and subjects were instructed to press, with fingers of the right hand, the left and right arrow keys to rotate the dial so as to replicate their memory of the orientation of the memorandum. The probe response window was 4 seconds. An inter-trial interval (ITI) of 3 seconds separated each trial.

The direction of the cue was randomly determined on each trial. The orientation of the memorandum was independently selected, with replacement, from a fixed set of 6 values spaced by 30 degrees (15°, 45°, 75°, 105°, 135°, and 165°) with a jitter between 0 to 3 degrees added. The congruity in orientation between memorandum and discriminandum was manipulated and counterbalanced across trials. On half of the trials the two orientations matched, whereas on the other half, they were either ±30, ±60, or 90 degrees apart. This manipulation resulted in 6 orientation values for the discriminandum, 15°, 45°, 75°, 105°, 135°, and 165°, the same as the memorandum values. The memorandum had a fixed contrast of 0.8. To enhance task engagement, the contrast of the discriminandum was varied and counterbalanced among 5 levels, (0.09, 0.07, 0.32, and 0.6). Noise was added to the discriminanda by replacing 30% of their pixels with random luminance. The orientations of the items in the uncued visual fields were also independently selected from the same pool of orientations. 24 subjects completed eight 60-trial blocks and one subject completed seven 60-trial blocks. Subjects completed the experiment in one session.

### Behavioral analysis

WM performance was measured in terms of precision and recall bias. Both within-trial and cross-trial effects were analyzed. Within-trial precision was determined by sorting trials based on the distance between the memorandum and discriminandum of trial *n*, and the inverse of the mean absolute error (precision =1/[mean absolute error]) was calculated for each distance condition. Cross-trial precision was determined by sorting trials based on the distance between the discriminandum of trial *n-1* and the memorandum of trial *n*.

To assess recall bias within trial, we first computed the mean signed error for each of the distance conditions between the trial *n* memorandum and the trial *n* discriminandum. A positive value would indicate an attractive bias of trial *n* recall toward trial *n*’s discriminandum, whereas a negative value would indicate a repulsive bias. Then we performed cross-trial serial dependence analyses with both model-free and model-based approaches. For the model-free approach, we calculated the median error (signed) for trials for which the relative distance between the discriminandum from trial *n* and the memorandum from trial *n+1* ranged from 0 to +/-45 degrees, and subtracted that from the median error on all trials within that range (e.g., Samaha et al., 2019; Teng et al., 2022).

For the model-based analysis of serial dependence we employed the derivative of gaussians procedure (Bliss, Sun, & D’Esposito, 2017; Fritsche, Mostert, & de Lange, 2017; Samaha, Switzky, & Postle, 2019; Teng et al., 2022): First, to prepare the data, trials with absolute error larger than 45 degrees were excluded (to minimize influences from random guesses). Next, we demeaned the data by subtracting the mean signed error across all trials from each trial’s error for each subject (in order to remove systematic response bias).

Subsequently, trials were sorted by the relative distance between trial *n-1* discriminandum and trial *n* memorandum. The data were smoothed by a 15-trial moving median filter, following previous conventions (e.g., Samaha et al., 2019; Teng et al., 2022). We then fit a derivative-of-Gaussian function (DoG; Fischer & Whitney, 2014; Bliss et al., 2017; Fritsche et al., 2017) to the data:

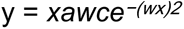

In which *x* is the relative orientation of trial *n-1* discriminandum, *a* corresponds to the amplitude of the peaks of the curve, *w* corresponds to the width of the curve, and *c* is a constant.

The DoG was fitted to the group-averaged data with two free parameters, amplitude (*a*; allowed to vary between -15 to 15 degrees) and width (*w*; allowed to vary between 0.02 to 0.2). A positive amplitude indicates an attractive serial bias; a negative amplitude indicates a repulsive serial bias. Width (*w*) scales the width of the tuning. The statistical significance of the DoG fits was determined with a bootstrapping procedure (10,000 interactions) in which we sampled 25 subjects with replacement and fit the DoG function to the group average data for each interaction. This procedure resulted in 10,000 values of the amplitude of the fit, and we compared this distribution against 0 to obtain the two-tailed p-value.

Performance on the discrimination task was assessed via response time (RT) and accuracy. Responses that were incorrect, or three standard deviations above or below the mean RT, were excluded from the RT analyses.

### EEG acquisition and preprocessing

Sixty three channels of EEG were recorded with BrainVision Recorder using active Ag/AgCl electrodes feeding into an ActiCHamp amplifier (Brainprodcuts GmbH, Gilching, Germany). The electrodes were placed in accordance with the extended international 10-20 system.

Impedances were kept below 25 kΩ. The data were sampled at 1000 Hz. An electrode placed on the forehead served as ground. All channels were referenced online to channel Fpz and re-referenced offline to the average of all electrodes.

Preprocessing and analysis of the data were completed in MATLAB (2021, The Mathworks) using custom routines and functions from EEGLab (Delorme & Makeig, 2004) and Fieldtrip (Oostenveld et al., 2010). The data were downsampled to 500 Hz and filtered offline with a band-pass filter (0.1 Hz - 100 Hz). Bad channels were identified by visual inspection and replaced via the spherical spline interpolation method. The data were epoched around the onset of the cue (-1000 ms to 1000 ms) and the onset of the discriminandum (-500 to 1000 ms).

Epochs with amplitudes above a threshold of ±100 μV were excluded from further analyses(M=3.9%; SD=3.17%). Artifacts related to eye blinks, horizontal eye movements, and muscles were corrected using independent component analysis (ICA) (M=2.6 components, SD=1.1 components).

#### Event-related potential (ERP)

For ERP analyses, an additional low pass filter (30 Hz, second-order Butterworth) was applied to the preprocessed data. The time window of interest was determined to be -200 to 800 ms around the onset of the discriminandum and the data were baseline corrected from -200 to 0 ms.

#### Time-frequency analysis

The preprocessed data were decomposed with a complex Morlet wavelet (1-35 Hz, 1 Hz frequency bins, 3-10 cycles; Cohen, 2019) with a custom routine in the Fieldtrip toolbox (Oostenveld et al., 2011). We then down-sampled the time-frequency results to 100 Hz to increase processing efficiency for subsequent analyses. The data were baseline-corrected to the average of all trial ITI from the window from -1000 ms to -800 ms relative to cue onset with the decibel conversion. (Note that, because of the anticipatory nature of proactive control, the time window of interest began prior to the trial onset, i.e., at -800 ms.

#### Flexible Control Model (FCM)

The levels of trial-wise proactive and reactive control were quantified with the FCM (Jiang et al., 2014; Jiang et al., 2015). The FCM uses a reinforcement learning algorithm with a flexible learning rate to account for the flexible adjustment of control in a changing environment, and provides trial-by-trial estimates of level of demand for proactive and for reactive control. It has been successfully applied to conflict adaptation tasks, such as a modified Stroop task (Jiang et al., 2014), revealing an insula-frontostriatal network involved in predicting and executing control (Jiang et al., 2015). It has also been used to elucidate the role of lateral prefrontal cortex in anticipatory control, via its application to data from an experiment applying transcranial magnetic stimulation to this region (Muhle-Karbe, Jiang, & Egner, 2018). Most recently, we have used the FCM to investigate WM-perception interactions on a behavioral task similar to the one presented here (Teng et al., 2022).

Details of the model have been described previously in Jiang et al. (2014, 2015), and its application a concurrent WM-perception tasks has been outlined in Teng et al., (2022). In summary, the model consists of three key parameters: predicted level of conflict (*f*), observed trial congruity (*o*), and a flexible learning rate (*α*). The observed trial congruity (*o*) is a binary variable (0-congruent; 1-incongruent) serving as the input to the model, while the model provides trial-wise estimates for the other two parameters. The model maintains a joint probabilistic distribution of the predicted conflict level (*f*) and the flexible learning rate (*α*). *f* represents the model’s anticipation of the probability of the forthcoming trial being incongruent (continuous, *f* ∈ [0, 1]), thereby operationalizing the level of demand for proactive control. The flexible learning rate (*α*) corresponds to the model’s belief in the volatility in the environment, and was not included in our analyses because volatility was not experimentally manipulated. For each trial *n*, the model has made a prediction of the level of conflict that will be present on that trial (*f_n_*) based on the joint distribution of *f* and *α*. As the trial gets underway, the congruity between memorandum and discriminandum (*o_n_)* is observed, and the distribution is updated based on a reinforcement learning rule: *f_n + 1_ ←f_n_ + α × (o_n_ − f_n_)*. Importantly, the “control prediction error” (*control PE*) corresponding to the difference between a trial’s predicted conflict *f_n_* and its subsequently observed congruity *o_n_* (i.e., *o_n_ − f_n_*) has two roles: quantifying the amount of reactive control required by the discriminandum (thereby triggering the recruitment of reactive control), and updating *f_n_* (for the remainder of trial_n_ and persisting across the ITI and into trial_n+1_ until it is updated by trial_n+1_’s discriminandum). On incongruent trials, a positive *control PE* is generated (between 0 to 1) and on congruent trials, a negative *control PE* is generated (between -1 to 0). For analyses that compared tertiles of trials of highest and lowest *control PE*, trials were selected separately for congruent and incongruent conditions. For incongruent trials, the tertile of trials with highest positive *control PE* values were selected, and for congruent trials, the tertile of trials with lowest negative *control PE* values were selected.

We performed a general linear model (GLM) analyses across time and space to investigate the neural correlates of reactive and proactive control in the EEG signals. For reactive control, to capture the phasic stimulus-evoked effects of the *control PE*, for each subject, we performed a multiple regression on the voltage data for all time points of interest within a window from -200 to 800 ms around discriminandum onset at each electrode. Regressors included *f*, magnitude of *control PE* (absolute value of *control PE*), and congruity. To characterize the hypothesized sustained effect of proactive control, we first calculated narrowband theta power by averaging across the frequency interval of 4-7 Hz. Then we performed the same type of regression analysis on theta power for each time point within a window from -1000 ms to 1000 ms around trial onset at each electrode, with *f* as the regressor of interest (combined with a constant term to form a GLM).

For the effects of reactive control in the discriminandum-locked ERP, significance testing was carried out on the regression weights of *f*, *control PE*, and congruity with comparisons against 0. For the effects of proactive control in theta power, we compared the regression weights of *f* against 0. Multiple comparisons were corrected for with a nonparametric cluster-based permutation test across time points and channels (10,000 permutations with the Monte Carlo method; Maris & Oostenveld, 2007). First, adjacent data points (one data point apart) and channels (a minimum of two required) with a *p*-value smaller than 0.05 (by two-tailed t-test) were clustered together. Then, we used the maxsum method (the sum of the *t*-values within each cluster) to determine the cluster-level statistic and comparing it to the maxsum of clusters identified in the null distribution obtained from the 10,000 permutations.

#### Multivariate decoding

Time-resolved multivariate decoding of the orientations of the memorandum and the discriminandum was performed on the voltage data of 17 posterior channels (P7, P5, P3, P1, Pz, P4, P6, P8, PO7, PO3, POz, PO4, PO8, O1, Oz, and O2), following Wolff et al. (2015, 2017). The preprocessed data were downsampled to 50 Hz to increase computational efficiency. We decoded orientation using Mahalanobis distance to compute the distances between the orientation on a given trial and the full range of possible orientations. Decoding was performed separately and independently for memorandum and discriminandum, and for items presented to the left and right of fixation within each participant. We used a leave-one-trial-out cross-validation method to calculate the decodability of the trial-wise orientation. For each time point of a given trial, its activity pattern was compared to each of the six orientation bins formed with the remainder of trials (-75°, -45°, -15°, 15°, 45°, 75°). The Mahalanobis distance between the left-out trial and each orientation bin was calculated with the covariance matrix from the training trials using a shrinkage estimator. The number of trials for each orientation bin was kept equal by randomly subsampling to match the smallest number of condition-specific trials. This procedure was repeated 100 times to obtain reliable estimates.

For the visualization of tuning curves, the pairwise distances were mean-centered and the sign was reversed. Thus, higher values indicate greater pattern similarity between a pair of orientations and lower values indicate less similarity between a pair. Decoding accuracy was calculated as the slope of the tuning curves. Lastly, for significance testing, cluster-based signed permutation tests were done in windows anchored to memorandum onset (0 to 800 ms) and to discriminandum onset (0 to 800 ms), with a cluster-forming and cluster-testing significance threshold of 0.05. Time-resolved decoding values were smoothed with a Gaussian kernel (3 time points) for visualization and statistical tests.

The decoding results were used to test two predictions about the effects of control on stimulus representation. First, to assess the possible influence of proactive control on enhancing the processing of the memorandum, we tested whether orientation decoding of the memorandum was higher for the tertile of trials with the highest *f* than the tertile of trials with the lowest *f* (permutation procedure; 10,000 permutations).

Second, to test the prediction that incongruent discriminanda with high *control PE*s would elicit reactive suppression, we compared the averaged decoding accuracy of the tertile of incongruent trials with the highest *control PE* versus the tertile of incongruent trials with the lowest *control PE* (each against zero, between the two; permutation procedure; 10,000 permutations with cluster correction). If unexpectedly incongruent discriminanda are indeed suppressed, their neural representation should be markedly different from those of the same stimuli on trials when they serve as congruent discriminanda. To examine this, we first constructed the representational profile for the memorandum by training a decoding model on the orientations of the memorandum. We then applied this model to decode the orientations of the discriminandum. During the testing phase, each test trial was labeled by the orientation of the discriminandum and we computed the neural distance between this test trial and the memorandum profiles. If a discriminandum and a memorandum share the same neural code, a discriminandum with a specific orientation (D1) would exhibit the closest distance to the memorandum of the same orientation (M1) relative to all other orientations (M2-M6) in memorandum space. Alternatively, if D1 is suppressed, its neural distance to M1 would be the furthest relative to all other stimuli M2-M6. Therefore, we predicted that incongruent discriminanda generating a high *control PE*s would reconstruct an inversed similarity pattern for that required additional involvement of reactive control. Similar approaches have been reported in previous studies (Yu, Teng, & Postle, 2020; Zhang & Luo, 2023).

## Results

### Behavioral results

#### Model-free analyses, visual discrimination

We first performed model-free analyses to investigate whether the congruity of trial *n* influenced visual discrimination performance on trial *n+1*. Repeated-measure ANOVAs with factors of trial *n* congruity and trial *n+1* congruity were conducted on discrimination RT and discrimination accuracy. For discrimination RT, we observed a significant main effect of trial *n* congruity, *F*(1, 24) = 4.68, *p* = 0.041, indicating that RT on trial *n+1* was faster following a congruent trial *n* compared to an incongruent trial *n*.

Neither the main effect of trial *n* congruity nor the trial *n* x trial *n+1* interaction reached significance *F*s < 0.26, *p*s > 0.61 (Figure 2.A). In terms of discrimination accuracy, the main effect of trial *n+1* congruity was significant, *F*(1, 24) = 8.00, *p* = 0.01, whereas the effect of trial *n* congruity was not, *F*(1, 24) < 0.001, *p* = 0.99. Furthermore, the congruity effect on trial *n+1* (the difference between congruent and incongruent condition) was reduced if trial *n* was incongruent compared to when it was incongruent, as indicated by a significant trial *n* x trial *n+1* interaction, *F*(1, 24) = 11.15, *p* = 0.003, a Gratton effect (Figure 2.A).

**Figure 2.**
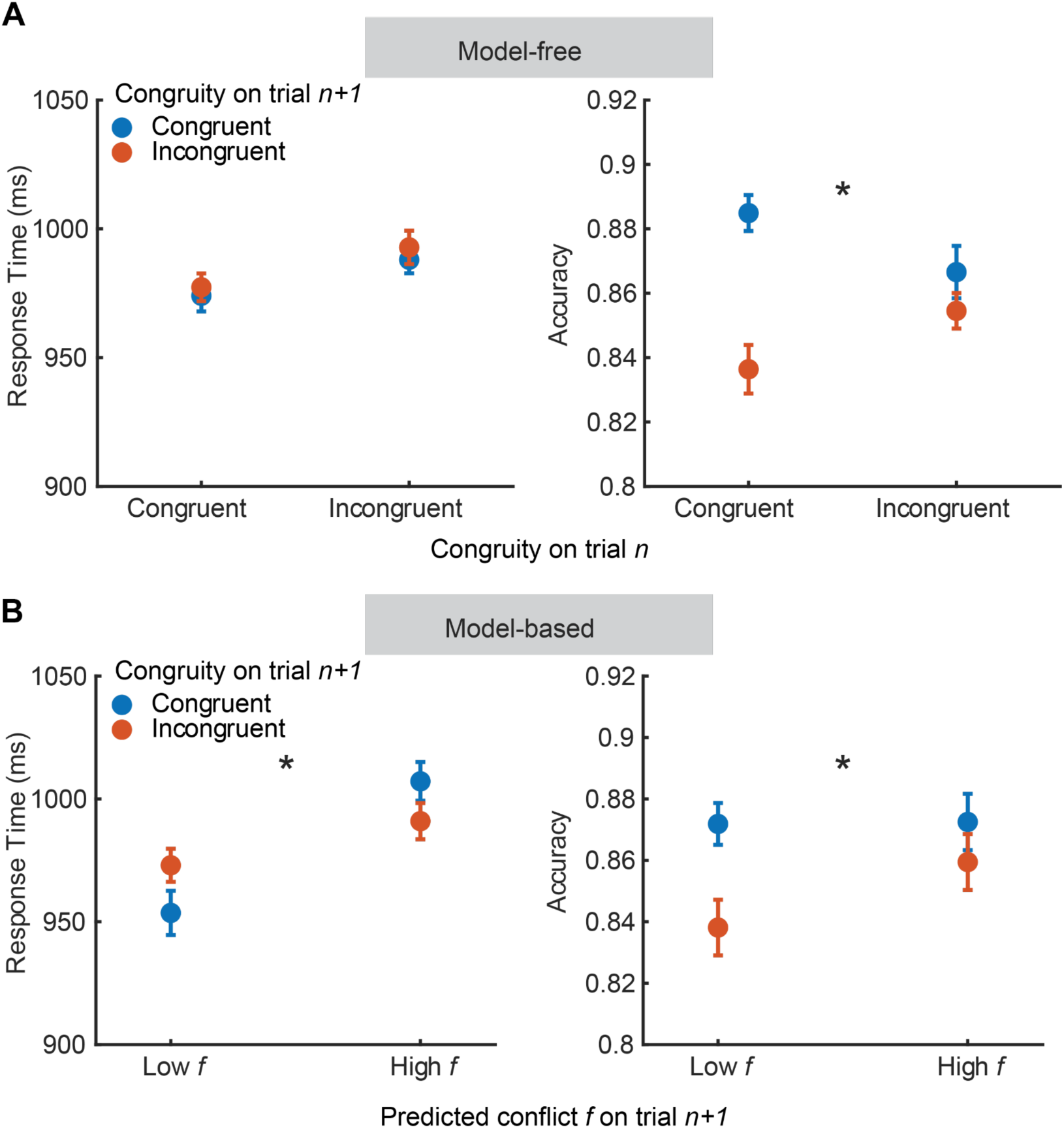
Behavioral effects of conflict adaptation on visual discrimination. (A) RT and accuracy on the discrimination task on trial *n+1*, as a function of trial *n* and trial *n+1* congruity. For accuracy, the congruity effect on trial *n+1* was reduced following an incongruent trial *n*: a reduction in both the interference caused by an incongruent discriminandum and the benefit caused by a congruent one, compared with following a congruent trial *n*. (B) Response time and accuracy replotted based on FCM estimates of predicted conflict *f* on trial *n+1*. *f* modulated the congruity effects for both RT and accuracy. * indicates *p* < 0.05 for the interactions; error bars indicate standard error (SE).

#### Model-free analyses, WM recall

Previous work with the dual-task procedure featured here (Teng et al., 2022), and with a WM retrocuing task (Shan and Postle, 2022), has provided evidence that when a stimulus elicits vigorous reactive control (e.g., an unexpected incongruent discriminandum) it can exert a repulsive serial bias, a reversal of sign of its influence when reactive control is not needed. To assess evidence for this phenomenon in the present experiment, trials were sorted by congruity, and a Difference of Gaussians (DoG) fitting of the bias was applied. Results indicated that although congruent discriminanda on trial *n* exerted the expected attractive bias on WM recall on trial *n+1* (*a* = 1.88°, *p* = 0.002, 95% CI = 0.94° to 2.70°, bootstrapped); incongruent discriminanda on trial *n* did not have a significant influence on WM recall on trial *n+1* (*a* = -0.61°, *p* = 0.10, 95% CI = -1.45° to 0.18°, bootstrapped). The amplitudes of the two curves were significantly different from each other (*p* < 0.001, bootstrapped).

Finally, precision in WM recall on trial *n+1* was influenced by trial *n+1* congruity, *F*(1, 24) = 11.56, *p* < 0.001, but not by trial *n* congruity, *F*(1, 24) = 3.78, *p* = 0.064, nor was the interaction significant, *F*(1, 24) = 1.09, *p* = 0.31.

#### Model-based assessment, proactive control

We operationalized the level of proactive control using the *f* parameter (predicted conflict) from the Flexible Control Model (FCM), which was found to modulate performance in visual discrimination. Trials were sorted by *f*, and the tertile of trial with the highest *f* and the tertile of trials with the lowest *f* were selected for further analyses. First, repeated-measures ANOVAs were conducted on discrimination RT and discrimination accuracy on trial *n+1*, with factors of *f* from trial *n+1* and congruity on trial *n+1*. (This has the effect of assessing the level of proactive control that was present at the onset of trial *n+1*.) For discrimination RT, we observed a significant main effect of *f*, *F*(1, 24) = 9.42, *p* = 0.005, and a significant interaction between *f* and congruity, *F*(1, 24) = 7.24, *p* = 0.013.

Specifically, trials that began with a higher *f* exhibited a reduced congruity effect (Figure 2.B). For discrimination accuracy, we found a main effect of congruity (*F*(1, 24) = 7.61, *p* = 0.011) and a significant *f* x congruity interaction (*F*(1, 24) = 4.36, *p* = 0.048) reflecting the fact that the congruity effect was absent in the presence of high *f* (Figure 2.B). For WM recall precision, only a significant main effect of trial *n+1* congruity was observed, *F*(1, 24) = 46.20, *p* < 0.001 (for the other effects, *F*s < 0.60, *p*s > 0.41).

#### Model-based assessment, reactive control

The model-free analyses indicated that, although the delayed recall and visual discrimination elements of the dual task are formally independent, they nonetheless interact, in that incongruity between memorandum and discriminandum worsens discrimination performance and WM recall precision (replicating Teng et al., 2022). It follows from this that the cognitive system may recruit reactive control in instances in which, when expected conflict (and, therefore, proactive control) is low, the discriminandum is (unexpectedly) incongruent with the memorandum. This intuition is better captured by the FCM than by the simple sorting of trials by congruity (see *Model-free analyses, WM recall,* above) because the *control PE* parameter factors is “the element of surprise”: It is instances of discrepancy between expected conflict and observed conflict that generate large control *PE*s. Indeed, a prediction arising from Teng et al. (2022), and tested here, is that the *control PE* recruits reactive control, in a manner such that the magnitude of the *control PE* predicts the amount of reactive control that is recruited.

For these analyses, trials were sorted by congruity and by the amount of the *control PE* parameter. Whereas congruent discriminanda eliciting high *control PE*s on trial *n* exerted the expected attractive bias on WM recall on trial *n+1* (*a* = 2.07°, *p* < 0.001, 95% CI = 0.96° to 3.25°, bootstrapped), incongruent discriminanda eliciting high *control PE*s on trial *n* exerted a repulsive bias on WM recall on trial *n+1* (*a* = -1.17°, *p* = 0.043, 95% CI = -2.42° to - 0.08°, bootstrapped), and the two curves differed significantly from each other (*p* = 0.002, bootstrapped; Figure 3.C). For the tertile of trials that were preceded by a trial with a congruent discriminandum that elicited a low *control PE*, the attractive serial bias exerted by the congruent discriminandum only trended toward significance (*a* = 1.47°, *p* = 0.058, 95% CI = 0.16° to 2.67°, bootstrapped), and for the tertile of trials that were preceded by a trial with an incongruent discriminandum that elicited a low *control PE* no serial bias was evident (*a* = -0.36°, *p* = 0.57, 95% CI = -1.28° to 0.68°, bootstrapped; Figure 3.C).

**Figure 3.**
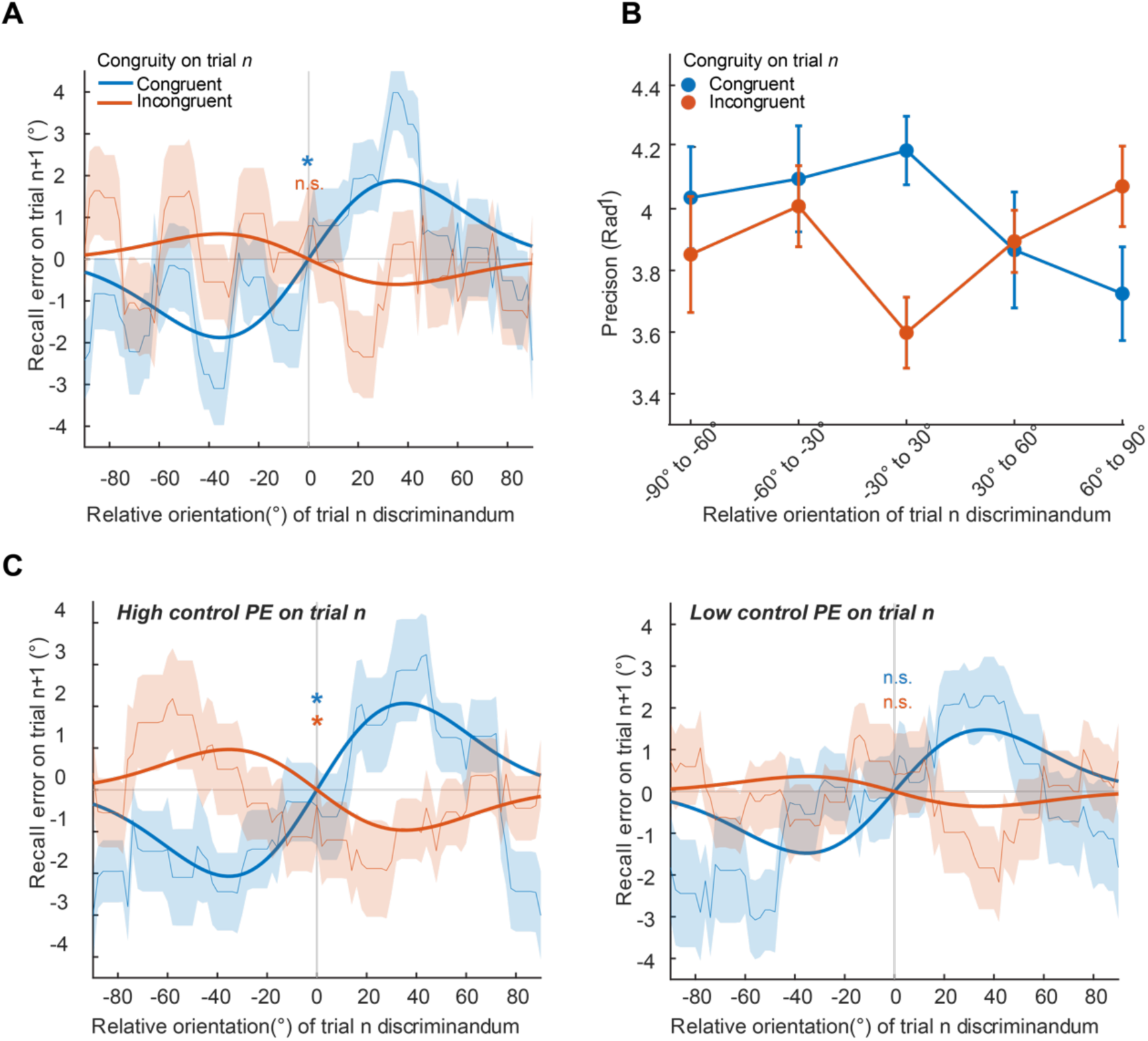
Behavioral effects of conflict adaptation on WM recall. (A) Serial dependence analysis of the influence of the trial *n* discriminandum on WM recall on trial *n+1*. After trial *n+1* data were sorted as trial *n*-congruent and trial *n*-incongruent, WM recall error on trial *n+1* was plotted based on the relative orientation of the trial *n* discriminandum. Only the attractive serial bias exerted by congruent discriminanda was statistically reliable. (B) WM recall precision on trial *n+1* plotted as a function of the relative angular distance from the discriminandum on trial *n*, revealing that orientation-dependent serial bias of the discriminandum was largely opposite as a function of congruity. (C) Influence of the trial *n* discriminandum on WM recall on trial *n+1* when trials are sorted by the *control PE* parameter from fits to the FCM. Left-hand panel: For the tertiles of congruent and incongruent discriminanda with the highest *control PE*s. congruent discriminanda exerted an attractive bias on WM recall on trial *n+1* and incongruent discriminanda exerted a repulsive bias on WM recall on trial *n+1*. Right hand panel: For the tertiles of congruent and incongruent discriminanda with the lowest *control PE*, *neither* congruent nor incongruent discriminanda exerted a significant serial bias. * indicates two-sided *p* < 0.05 against 0.

Results of the analysis of WM recall precision on trial *n+1* as a function of the relative angular distance from the discriminandum on trial *n* (Figure 3.B) are consistent with the possibility that the reactive control triggered by an incongruent discriminandum results in feature-specific suppression of the representation of that discriminandum. Previous work has documented impairments in performance when a suppressed feature reappears as a visual search target (e.g., Sawaki & Lick, 2010; Gaspelin et al., 2015) or a WM sample (Teng et al., 2022).

Therefore, we examined whether the precision of WM recall was modulated by the congruity of the discriminandum from the preceding trial in an orientation-dependent manner. To do this we first sorted trials by congruity on trial *n*, then further divided them into five bins according to the relative orientation difference between the discriminandum from trial *n* and the memorandum from trial *n+1*: -90° to -60°, -60° to -30°, 0° to 30°, 30° to 60°, and 60° to 90° (Figure 3A (right). For statistical analysis negative orientation distances were folded over to the positive side to increase power, resulting in three bins: 0° to ±30°, ±(30° to 60°), and ±(60° to 90°). A repeated measure ANOVA with factors of trial *n* congruity and orientation distance revealed a significant interaction between the two factors, *F*(2, 48) = 6.04, *p* = 0.005. Post Hoc comparisons indicated that when a congruent discriminandum was followed by a memorandum with a similar orientation (within the 0° to ±30° range) on the next trial *n+1*, the recall precision of that memorandum was enhanced (*p* = 0.04 compared with ±[30° to 60°]; *p* = 0.019 compared with ±[60° to 90°], Bonferroni corrected).For an incongruent discriminandum, in contrast, there was a trend suggesting that recall precision was worse when the memorandum from trial *n+1* was similar in orientation to the discriminandum from trial *n*, although none of the pairwise comparisons survived Bonferroni correction. Note that we did not perform this orientation-dependent analysis in relation to the FCM parameter estimates due to the limited number of trials per bin (less than 20), which would yield unreliable estimates of recall precision.

### EEG results

#### Proactive control elevates pre-trial theta-band power at frontal midline electrodes and enhances WM representation

Within the framework of the FCM, the level of proactive control (*f*) during performance of the WM+discrimination dual task is updated on each trial by the *control PE* elicited by the discriminandum on trial *n*, and this new level of *f* persists until it is updated by the *control PE* on trial *n+1.* Thus, it should manifest as a tonic signal that persists from shortly after the processing of the discriminandum on trial *n* until it is updated by the *control PE* triggered by the discriminandum on trial *n+1*. To search for a neural correlate of such a signal, we focused on theta-band oscillations within a time window of interest defined as -800 ms to 1000 ms relative to trial onset. The data were baseline-corrected from -1000 ms to -800 ms relative to trial onset, and theta-band power was extracted as the 4-7 Hz band passed signal from all electrodes. The time series signal was regressed against *f*, and the resultant regressor weights run through a cluster-based permutation test in 2D (channel x time point). This analysis revealed a cluster of electrodes along the frontal midline that exhibited significant regression weights for *f*. The significant loading on frontal midline theta started to emerge around -460 ms and lasted until -40 ms relative to the onset of the trial (Figure 4A-B, *p* = 0.03). Figure 4C illustrates the spectral difference between the tertile of trials with the highest *f* and the tertile of trials with the lowest *f* at electrode Cz.

**Figure 4.**
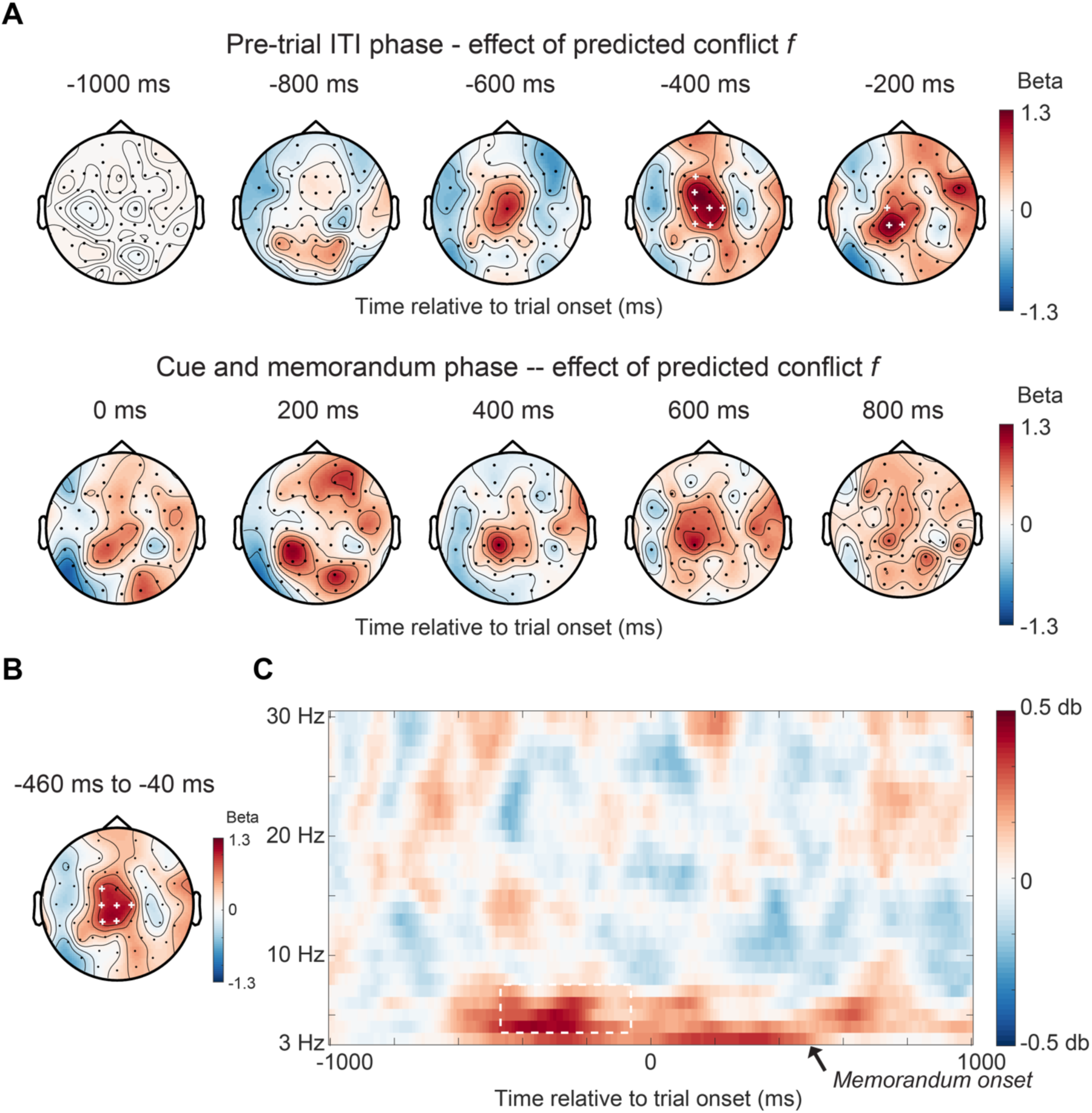
Neural signature of proactive control – pretrial theta power at frontal midline electrodes scales with the level of expected conflict. (A) Scalp topography of regression weights for the *f* parameter from the FCM derived from regression analysis on theta-band (4-7 Hz) activity. Top row: 200 msec windows of the final second of ITI before onset of trial *n+1* (with *f* having been updated by the discriminandum-triggered *control PE* from trial *n*); bottom row: 200 msec windows of the first sec of trial *n+1*. (B) Topography of the cluster of electrodes with significant correlation of *f* with theta-band power between -460 ms and -40 ms relative to trial onset. (C) Time-frequency plot at electrodes from the identified cluster (Cz, C1, C2, FC1, CP1, and CPz) showing an increase of theta-band power before the onset of the trial. The white dashed outline indicates the significant time points from the cluster in (B) identified through the cluster-based permutation test.

To investigate how the level of proactive control may influence WM processing, we conducted a multivariate decoding analysis of the representation of the orientation of the memorandum. On trials with high *f* reliable decoding of the memorandum emerged roughly 100 ms after stimulus onset (*p* = 0.038, cluster corrected) On trials with low *f*, however, above-baseline decoding did not emerge until roughly 400 ms following stimulus onset (*p* = 0.050, cluster corrected). Furthermore, decoding accuracy was significantly higher on high *f* trials compared to low *f* trials during the early stages of memorandum presentation (80-100 ms after memorandum onset, *p* = 0.039, cluster corrected, Figure 5B and 5C).

**Figure 5.**
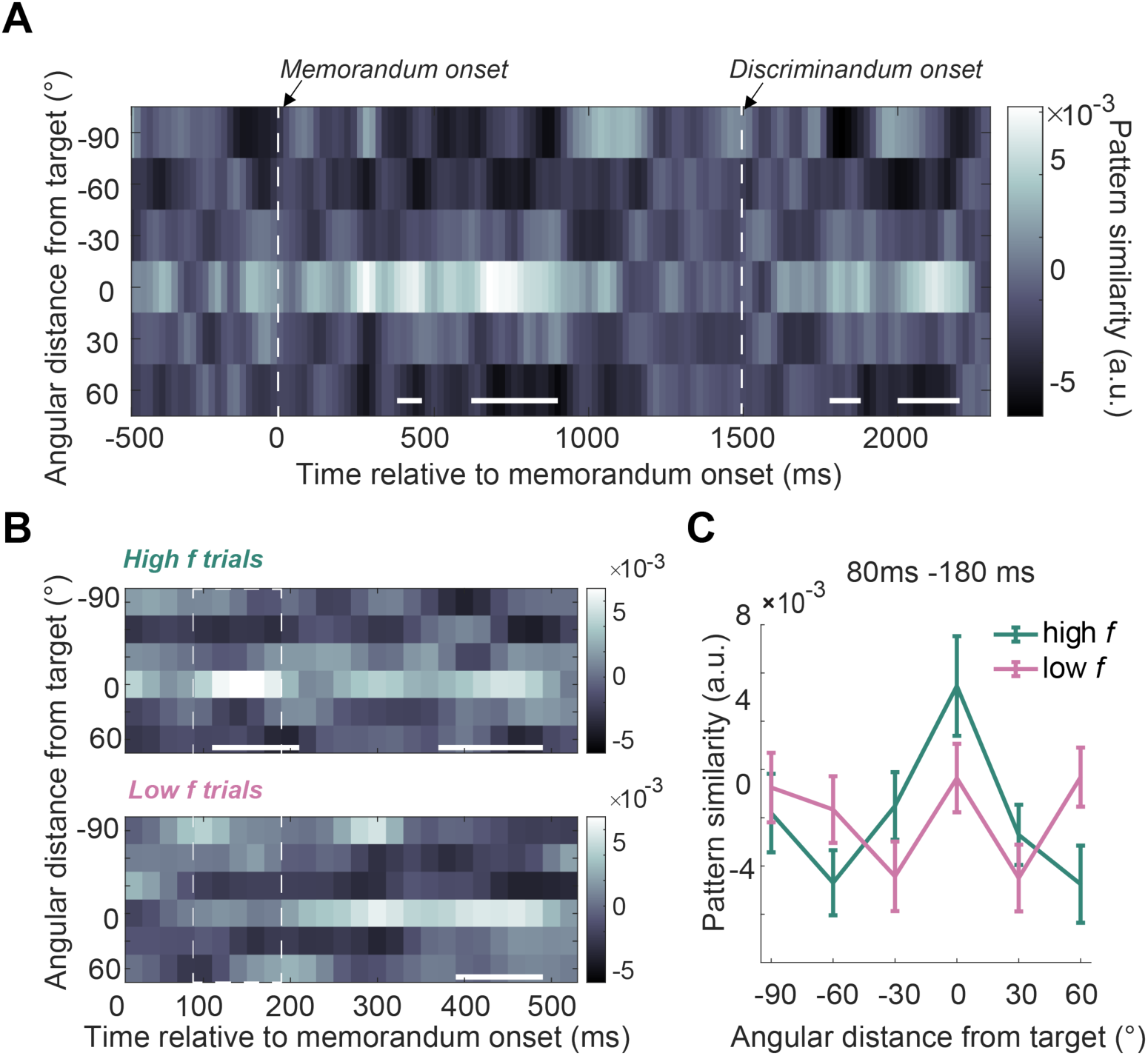
Influence of proactive control on the neural representation of the memorandum. (A) Time course of memorandum decoding averaged across all trials. Similarity patterns (Mahalanobis distance, sign-reversed) from a single test trial to the training template were mean-centered and averaged for each angular distance and time point. Decoding was successful after the onset of the memorandum and the onset of the discriminandum. Solid white lines mark statistically significant time periods (two-tailed p < 0.05, cluster corrected). (B) Average similarity patterns for memorandum orientation on high *f* and low *f* trials. Whereas on high *f* trials the memorandum was quickly decodable after its onset, on low *f* trials it was not decodable until around 400 ms. Solid white lines indicate statistically significant time periods (against zero; p < 0.05, cluster corrected). The dashed white lines mark time points at which high and low f trials are statistically different. (C) Similarity patterns averaged over 80-180 ms identified in (B) illustrate more robust stimulus representation on high *f* trials than low *f* trials.

#### The interplay between proactive and reactive control during perceptual discrimination

Within the framework of the FCM, discriminandum onset generates the binary value of observed trial congruity (*o*), and the difference between o and *f* determines the sign and amplitude of the resultant *control PE* which, in turn, triggers the engagement of the reactive control that may be needed to control the consequences of memorandum-discriminandum congruity. Because the *control PE* is a transient signal, we focused on discriminandum-locked ERPs to examine these dynamics (-200 ms to 1000 ms relative to discriminandum onset; baseline-corrected to 200 ms prior to discriminandum onset). Regression analyses were performed with *f*, *control PE*, and trial congruity as regressors against EEG voltage (i.e., the ERP), revealing distinct timing and scalp topographies for each regressor (Figure 6). Beginning with *f*, its influence emerged early (152 ms to 256 ms after discriminandum onset, *p* = 0.03, cluster corrected), with a significant cluster forming around the central electrodes that showed increased positivity with higher values of *f* (Figure 6.A). The influence of *o* was seen at posterior electrodes, with incongruent trials displaying increased negativity from 444 ms to 800 ms (*p* = 0.02, cluster corrected; Figure 6.B). Finally the *control PE* elicited a left frontal positivity from 420-742 ms post discriminandum onset (*p* = 0.04, cluster corrected, Figure 6C), thereby appearing at a similar time as the effect of *o*.

**Figure 6.**
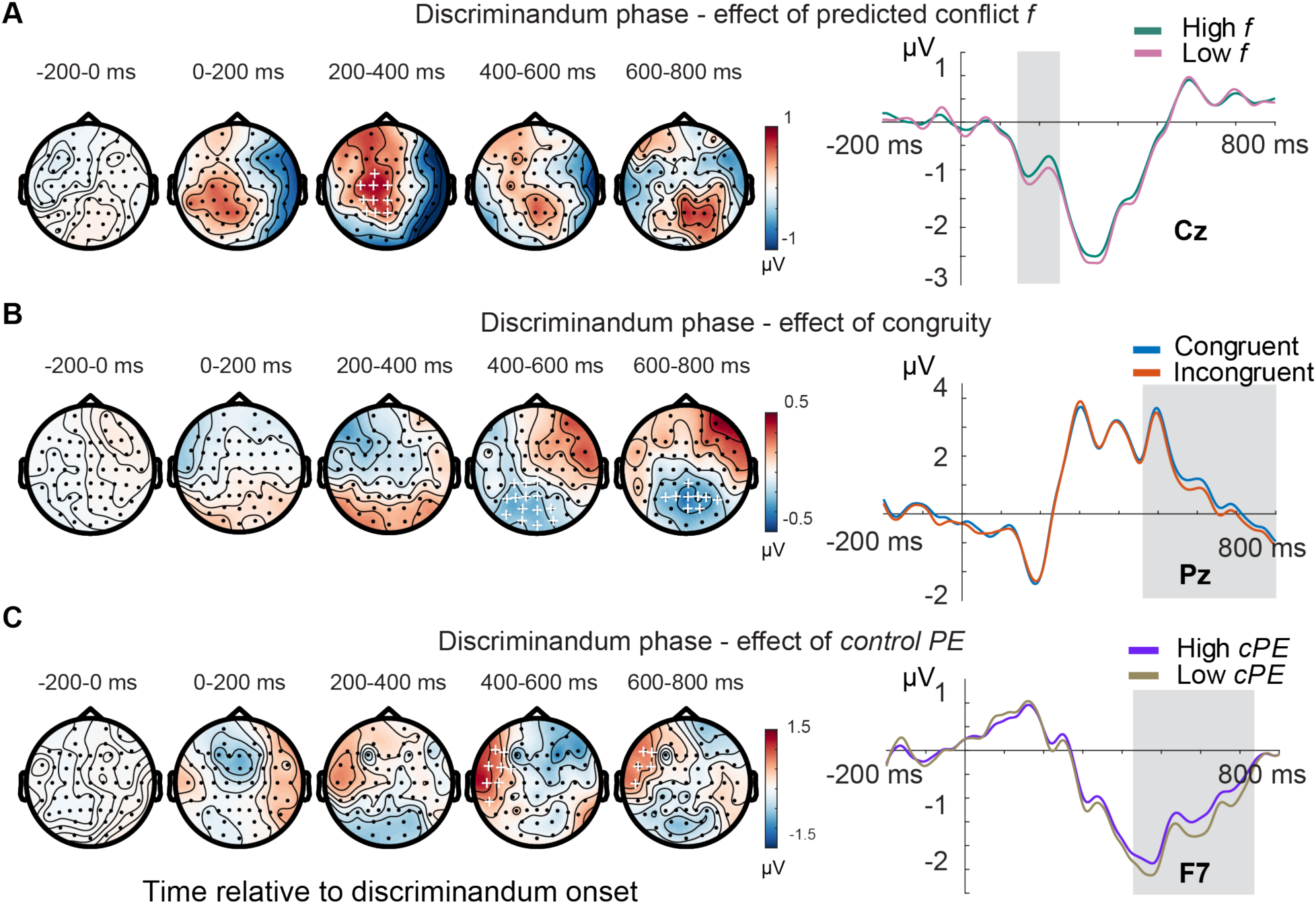
The interplay between proactive and reactive control during the discrimination task. (A) Left: scalp topography of regression weight for predicted conflict (*f*). Discriminandum-locked ERP from a large cluster of central electrodes was sensitive to *f* from ∼150-250 ms. Right: ERPs plotted from Cz, comprised of trials with the highest and lowest tertile of values of *f*. (B) Left: scalp topography of regression weight for congruity between the memorandum and the discriminandum (*o*), revealing a cluster of posterior electrodes that was modulated by congruity beginning at ∼450 ms after discriminandum onset. Right: ERPs plotted from Pz for congruent and incongruent trials. (C) Left: scalp topography of regression weights for *control PE*, revealing a cluster of left-frontal electrodes that were significantly modulated by *control PE* from ∼420-742 ms following discriminandum onset. Right: ERPs plotted from F7 for the highest and lowest tertile of trials for *control PE*. Gray shaded areas indicate time points of significant difference identified by the regression analysis (two-tailed *p* < 0.05, cluster-corrected). The timing of these effects is consistent with the temporal dynamics hypothesized by the FCM: an early engagement of proactive control (*f*; because it was present at this level prior to discriminandum onset), and subsequent comparison of observed conflict (*o*) against *f* to generate the *control PE*.

Figure 6D illustrates the difference between the tertile of trials with the highest and lowest *f*, congruent and incongruent trials, and the tertile of trials with the highest and lowest *control PE*. These results were consistent with the hypothesis that proactive control is engaged during the early processing of the discriminandum, and the *control PE* emerges upon the processing of discriminandum congruity.

#### Neural consequences of reactive control

The behavioral data from this study and from Teng et al. (2022) indicate that an incongruent discriminandum impairs recall of the memorandum at the end of the trial. We have previously hypothesized that the reactive control triggered by an unexpectedly incongruent discriminandum has the effect of suppressing its neural representation, so as to minimize its potential to disrupt WM recall (Teng et al., 2022). One of the behavioral hallmarks of such suppression may be the repulsive serial bias exerted by incongruent discriminanda that generate high *control PE*s (Fig. 3; Teng et al., 2022), an effect that has also been observed for items that are actively removed from WM (Shan and Postle, 2022). Here, we assessed neural evidence for suppression-by-reactive control by examining the neural coding of discriminanda in different conditions.

Figure 7A showed the similarity between the neural representations of the discriminandum and the memorandum. A congruent discriminandum with a high *control PE* could be reliably decoded from the model trained on memorandum orientations (280 ms to 480 relative to discriminandum onset, two-sided *p* = 0.035, cluster-corrected). Critically, the incongruent discriminandum with a high *control PE* was associated with a “flipped” representation where the similarity pattern had the lowest value at the target orientation (360 ms to 460 relative to discriminandum onset, two-sided *p* = 0.021, cluster-corrected), suggesting that its neural representation was flipped relative to its representation when a memorandum. The congruent and incongruent discriminanda significantly differed from each other during the discriminandum-locked time window of 200 ms to 480 ms. For low *control PE* trials, a brief period of decodability of the congruent discriminandum was observed at a later time point with a positive tuning profile (640 ms to 700 ms, two-sided *p* = 0.048, cluster-corrected). Figure 7B depicts the tuning profiles for those four trial types during the time window when congruent and incongruent discriminanda on high *control PE* trials differed significantly (200 to 480 ms). These findings align with the behavioral serial dependence results, and provide direct neural evidence for the active involvement of reactive control, whereby incongruent perceptual input is actively suppressed to resolve conflicts between it and the contents of working memory.

**Figure 7.**
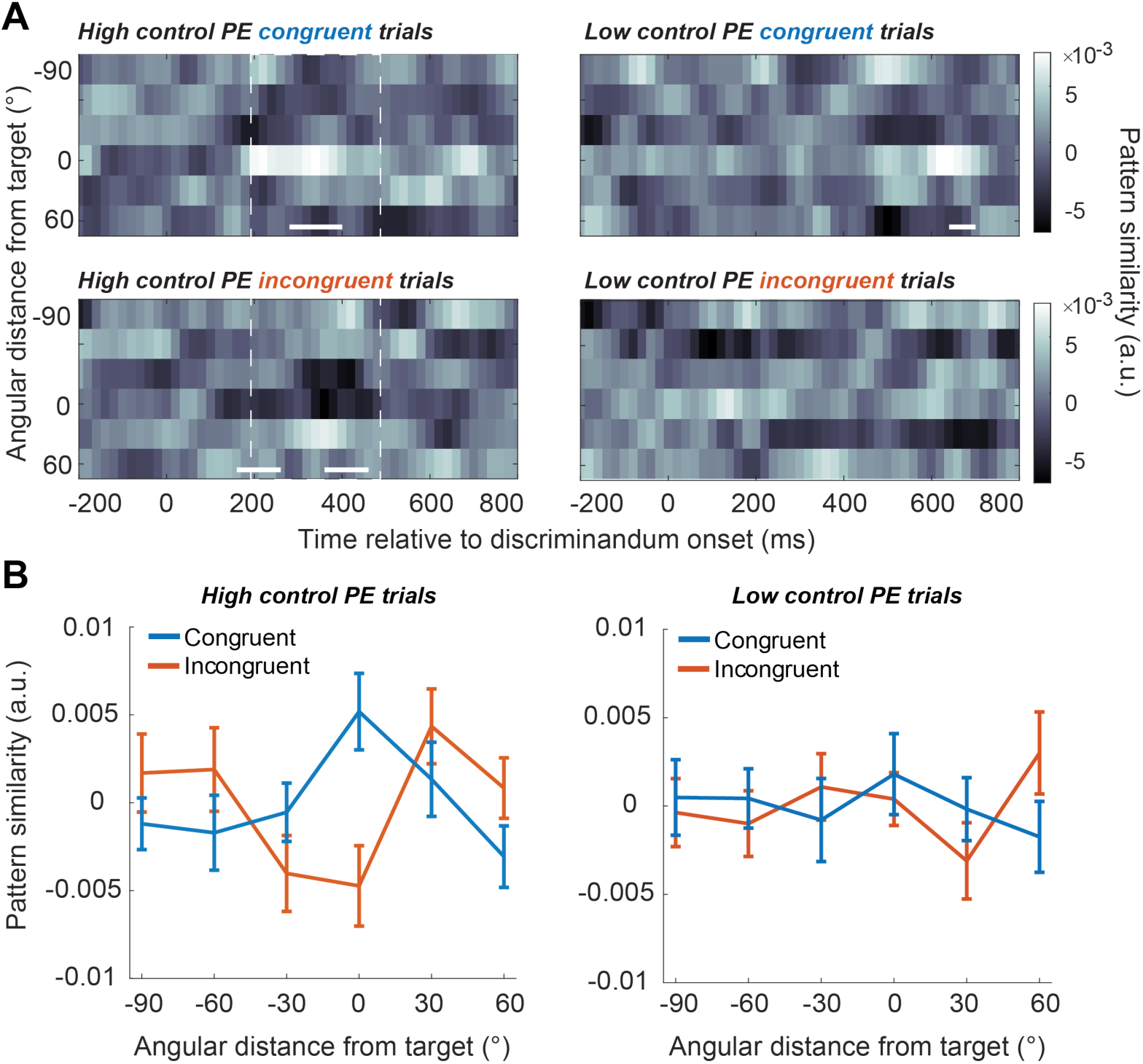
Neural evidence of reactive suppression as a consequence of control predictor error. (A) Time-resolved decoding results for the discriminandum. For high *control PE* trials, congruent discriminanda were represented in the same active format as the memoranda, whereas incongruent discriminanda were represented in a reversed format, consistent with suppression. Solid white lines indicate significant decoding (two-sided p < 0.005, cluster-corrected), dashed white lines demarcate the span of time when the two differ significantly (i.e., 200-480 ms relative to discriminandum onset). (B) Averaged similarity patterns between 200 to 480 ms for the four trial types.

## Discussion

In this study, we investigated the role of two modes of cognitive control in mitigating interference between perception and WM. Although previous research has demonstrated the engagement of control during WM-perception interactions, here we used model-based EEG to track the temporal dynamics of these hypothetically distinct mechanisms, as well their influence on perceptual and mnemonic representations.

Our study successfully applied the Flexible Control Model (FCM), originally developed for conflict adaptation tasks that lack an explicit demand on WM (e.g., the Stroop task, to tease apart proactive from reactive control on a trial-by-trial basis. After establishing that our behavioral results replicated the findings from previous work (Teng et al., 2022) -- that parameter estimates from the FCM explain key effects of the control of conflict on performance on the dual WM-plus-discrimination task – we directly linked these parameter estimates to EEG signals via multiple-regression analysis. First, we observed that theta-band power at frontal midline electrodes closely tracked the level of proactive control, a signal that, updated by recent trial history, was most prominent prior to trial onset. With the onset of the trial, the effects of proactive control were evident in the neural reconstruction (i.e., the representational strength) of the memorandum, and in early components of the discriminandum-locked ERP. Furthermore, EEG components tracking the effect of congruity and the *control PE* emerged around the same time, suggesting dynamic interactions that give rise to reactive control. Although context-dependent shifts between proactive and reactive control have been broadly established, studies have typically manipulated conflict either at the list-wise or at the item-specific level in order to create task contexts that favor one mode of control over the other (e.g., Braver et al., 2009; Braver et al., 2021; Bugg et al., 2013; Gonthier et al., 2016a; Gonthier et al., 2016b). To our knowledge, there has been only limited behavioral evidence to date that examines the simultaneous engagement of the two modes of control (Mäki-Marttunen et al., 2019). The results presented here provide a novel demonstration of the concurrent recruitment of proactive and reactive control at the single trial level, and illustrate how these two modes of control interact, engaging distinct neural processes to minimize conflict-induced interference.

The finding linking theta-band power at frontal midline electrodes with the level of proactive control is consistent with the broader literature on cognitive control, and one influential view is that “frontal-midline theta” (FMθ) represents the general need for, and anticipation of the need for, control (e.g., Cavanagh et al., 2012; Cavanagh & Frank, 2014; Chang et al., 2017; Cooper et al., 2019; Kaiser et al., 2019; Messel et al., 2021). For example, one recent study showed that in a conflict action paradigm, presenting a predictive cue to indicate upcoming conflict prior to the action signal led to increase in FMθ power during the preparation phase, indicating a role in conflict preparation (Messel et al., 2021). Here, our results showed that trial-by-trial levels of FMθ power fluctuated with predicted need for the control of conflict in an anticipatory fashion, prior to trial onset, thus providing additional support for the link between FMθ and anticipatory control. Furthermore, FMθ has been studied primarily in task contexts involving response conflict or response inhibition (e.g., Cavanagh et al., 2012; Chang et al., 2017; Messel et al., 2021). In the present study, incongruity occurred at the level of stimulus representation, putting the loci of conflict not on the response, but rather on the processes of perceptual decision making and WM recall. Our findings thus suggest that FMθ signalling may serve as a general mechanism for regulating top-down bias on sensory and motor processing networks, even in the absence of overt response conflict. Interestingly, one of the EEG results presented here has also provided evidence for a neural effect of proactive control for which we have not found a behavioral correlate. This relates to the fact that although WM recall accuracy suffers on incongruent trials, this effect does not seem to be mediated by control (model-free: it is insensitive to congruity on trial n-1; model-based, it is insensitive to *f*). Nonetheless, decoding of the memorandum is stronger at high levels of *f*, an effect that would typically be interpreted as evidence for stronger encoding into WM. Although this requires further study, it is possible that any control-related enhancement of encoding into WM in this WM-plus-discrimination task is diluted by the events that intervene between encoding and recall.

Turning to the *control PE*, our analyses identified a cluster of left anterior electrodes for which voltage scaled positively with the magnitude of the *control PE*. This effect is consistent with past research that has implicated lateral PFC in conflict resolution (e.g., Burgess & Braver, 2010; Feredoes et al., 2006) and in learning-dependent control (Jiang et al., 2015). In a color naming Stroop task that compared the coding schemes in DLPFC and dorsomedial frontal cortex (DMFC), DMFC was found to encode the trial incongruity (control demand) and DLPFC both target and incongruity-related information. Importantly, stronger targe coding in DLPFC was associated with a reduced Stroop interference (Freund et al., 2021), consistent with the proposed functionally dissociated roles of DMFC in control demand monitoring and of DLPFC in control implementation (Braver & Cohen, 2000; Shenhav et al., 2013). For WM, conflict processing has been studied extensively with a ‘recent-negative probes’ procedure that manipulates proactive interference from previously encountered information (e.g., Jonides et al., 1998; Jonides & Nee 2006;Monsell, 1978). The common finding with this procedure is that a probe that was not been a member of the current trial’s memory set but was a member of the previous trial’s memory set takes longer to reject and elicits increased activity in left inferior frontal gyrus (IFG; e.g., Jonides et al., 1998; Postle et al., 2004). Furthermore, repetitive transcranial magnetic stimulation (rTMS) applied to left IFG concurrent with probe onset selectively exaggerates the RT cost associated with recent-negative probes (Feredoes et al., 2006; Feredoes and Postle, 2009).

In the context of the FCM, model-based analyses of neuroimaging data have shown that, in a modified Stroop task, activity in left anterior insula and adjacent IFG is linked both to the magnitude of the *control PE* and to the flexible learning rate in the FCM that governs the amplitude of the change to *f* that is produced by a given value of the *control PE* (i.e., the α parameter from the FCM, which we did not manipulate in the present study). environmental volatility (flexible learning rate) (Jiang et al., 2015). Furthermore, rTMS of left lateral PFC eliminates behavioral effects of phasic and tonic adaptation to changes in the probability of incongruent trials, an effect interpreted as a disruption of the processes by which the *control PE* drives updates to *f* (phasic adaptation) and to α (tonic adaptation; Muhle-Karbe et al., 2018). Thus, the present results are consistent with the idea that the left lateral PFC contributes to the processing of the *control PE* that leads to flexible adjustments in the control of proactive control.

The present results also provide novel evidence about mechanisms of reactive control. Recall that the need for reactive control of incongruent discriminanda is illustrated by the fact that WM recall suffers as the angular difference between a trial’s memorandum and the previous trial’s discriminandum decreases (Figure 3.b). Behaviorally, the fact that discriminanda that generate large *control PE*s also exert stronger serial biases (Fig., 3, c) has been interpreted as evidence for a role for the *control PE* in triggering reactive control (Teng et al., 2022). Here, analyses of the concurrently recorded EEG data provide neural evidence for this link: Multivariate decoding of discriminanda from the EEG data reveals that, on high *control PE* trials, the neural representations of incongruent discriminanda are suppressed below baseline, a presumed consequence of reactive control. This finding suggests a neural explanation for why the serial bias exerted by an item from trial *n* on WM recall on trial *n+1* will be either repulsive or attractive as a function of whether or not its representation needs to be suppressed (Shan and Postle, 2022; Teng et al., 2022). It also supports the interesting possibility that active suppression may be a common mechanism that the visual system employs to reduce interference from task-irrelevant information in many contexts, including visual perception (e.g., Gaspelin et al., 2015) and visual WM (Shan & Postle, 2022; Teng et al., 2022). Although model-based analyses implicate the *control PE* as a signal that can trigger reactive control, understanding the source(s) of the reactive control signal itself will require further work.

To conclude, our results demonstrate the trial-by-trial neural dynamics associated with the interplay between the adaptive adjustment of proactive control and the recruitment of reactive control in the control interference between WM and perception. In addition to the control signals themselves, they also characterized the consequences of these control processes for the neural representation of stimulus information in the visual system. Understanding the neural dynamics of distinct control mechanisms for conflict resolution is essential for gaining insights into flexible and adaptive human behavior.

